# Leukemia confers a durable imprint on healthy hematopoietic stem and progenitor cells

**DOI:** 10.1101/2024.11.01.621509

**Authors:** Ding-Wen Chen, Julie M. Schrey, Jian-Meng Fan, Sarah E. Adams, Deanne M. Taylor, Eric K. Wafula, Peter Kurre

## Abstract

Recent models of infection and experimental inflammation reveal that hematopoietic stem and progenitor cells (HSPCs) can generate a memory of the exposure that heightens the response to subsequent stimulation, a process termed central trained immunity. Inflammation is also a constitutive feature of cancer, including hematologic malignancies. Here, we adapt a translationally relevant model of acute myeloid leukemia (AML) to determine if inflammation in the bone marrow (BM) niche durably reprograms resident healthy HSPCs. To simulate the onset of malignancy along with the associated inflammatory surge as well as the subsequent remission, we generated hematopoietic chimera with healthy HSPCs and HSPCs bearing a doxycycline-responsive oncogene (hMLL-AF9) expression cassette, a validated model of AML. Results show that the exposure to AML blasts in the BM leaves healthy HSPCs during experimental remission with broad transcriptomic, epigenetic changes and enhanced reliance on glycolysis. A heterologous secondary challenge of AML-experienced animals resulted in pronounced gene expression changes in inflammatory and metabolic pathways. These augmented responses coincided with altered chromatin accessibility in AML-experienced HSPCs. Motif analysis of the epigenome in AML trained HSPC points to the involvement of core hematopoietic transcription factors. Altogether, these observations provide first evidence for the durable inflammatory reprogramming of healthy HSPCs in the cancer microenvironment.

## Introduction

Inflammation describes a complex tissue response that acts as an evolutionarily conserved, first line of defense against non-self signals. Conveyed through coordinate action of adaptive and innate immune effector cells, inflammation is inherently self-limited and resolves with removal of the stimulus. Chronic inflammation, by contrast, is a pathophysiological outcome that compromises tissue function, with systemic sequelae for hematopoietic, cardiovascular, hepatic and renal function^1^. Understanding the triggers and sustaining features of inflammation is a critical first step to address excess the associated morbidity and mortality. Studies over the past decade have defined a central role for hematopoietic cells in systemically propagating inflammation^2^. Examples of experimental autoimmunity, periodontitis and osteoarthritis illustrate the systemic inflammatory crosstalk that leads to engagement of the hematopoietic stem and progenitor cell (HSPC) compartment and their mature myeloid progeny central to this process^3–5^. Chronic periodontitis, for example, fuels a self-reinforcing process with expansion of clonal HSPC populations that bear recurrent risk mutations in the bone marrow, a condition termed clonal hematopoiesis of indeterminate potential (CHIP)^6^. CHIP in turn leads to well documented inflammatory co-morbidities in older individuals^2^. Like earlier studies of experimental infection or sterile inflammation, the underlying mechanism involves an epigenetic imprint generated by the original inflammatory exposure with metabolic reprogramming and transcriptional hyperresponsiveness to a subsequent inflammatory challenge. Together, these are now considered defining features of a process termed central Trained Immunity (cTI)^7^.

Inflammation is also a hallmark feature of cancer^8^. For example, in patients with acute myeloid leukemia (AML), leukemic blasts and bone marrow stroma contribute to an inflammatory bone marrow (BM) secretome that suppresses healthy hematopoiesis^9,10^. The profound impact on patient outcomes is evident in a recent study that identified inflammation as an independent risk factor in prognosis and survival of AML patients across age groups and leukemic subtypes^11^. HSPCs are components of the innate immune system that sense and secrete a broad range of inflammatory factors and overwhelmingly predominate in the BM once they re-expand after leukemic remission. Whether cancer can confer cTI on HSPC is not known, but we reasoned that such an overarching impact of inflammation on AML outcomes, well beyond remission, may reflect a reprogrammed hematopoietic compartment as a durable source of inflammation.

We recently showed that the inflammatory environment initiated by the expansion of AML blasts in the BM niche actively recruits long-lived HSPCs^12^. The present study aimed to address if persisting “AML-Experienced” (AML^EXP^)- HSPC acquire and retain features of cTI after disease remission.

Results in a chimeric model of doxycycline (DOX)-inducible human MLL-AF9 (MLL-AF9) AML show that leukemia initiation and sequential regression leave AML^EXP^-HSPC with epigenetic and metabolic adaptations that support an amplified response to a secondary inflammatory challenge, consistent with a memory of cancer in long-lived HSPCs.

## Results

### Generation of a chimeric model to simulate AML remission

Previously, we found that AML, even at low leukemic burden, leads to inflammatory activation of healthy HSPCs in the BM niche^12^. We reasoned that while hematopoietic function returns to steady state in remission, sterile AML-mediated inflammation may have durable consequences in HSPCs. To model the “AML-experienced” (AML^EXP^) remission state in HSPCs, we generated inducible AML chimera (AF9^+/+^) by co-transplanting wildtype (WT; CD45.1/2+) and inducible hMLL-AF9 (CD45.1+; homozygote hMLL-AF9 and rtTA) whole BM at a 1:4 ratio into myeloablated C57BL/6J recipients. AF9^+/+^ cells express a truncated nerve growth factor (NGFR) epitope for convenient detection^13^. Mice transplanted with WT and hMLL-AF9-null (CD45.1+; homozygote rtTA) BM served as controls (AF9^-/-^). Once engrafted to steady state after 6-8 weeks, AML was induced in AF9^+/+^ mice via DOX as previously described (Figure 1A)^12,14^.

**Figure 1.**
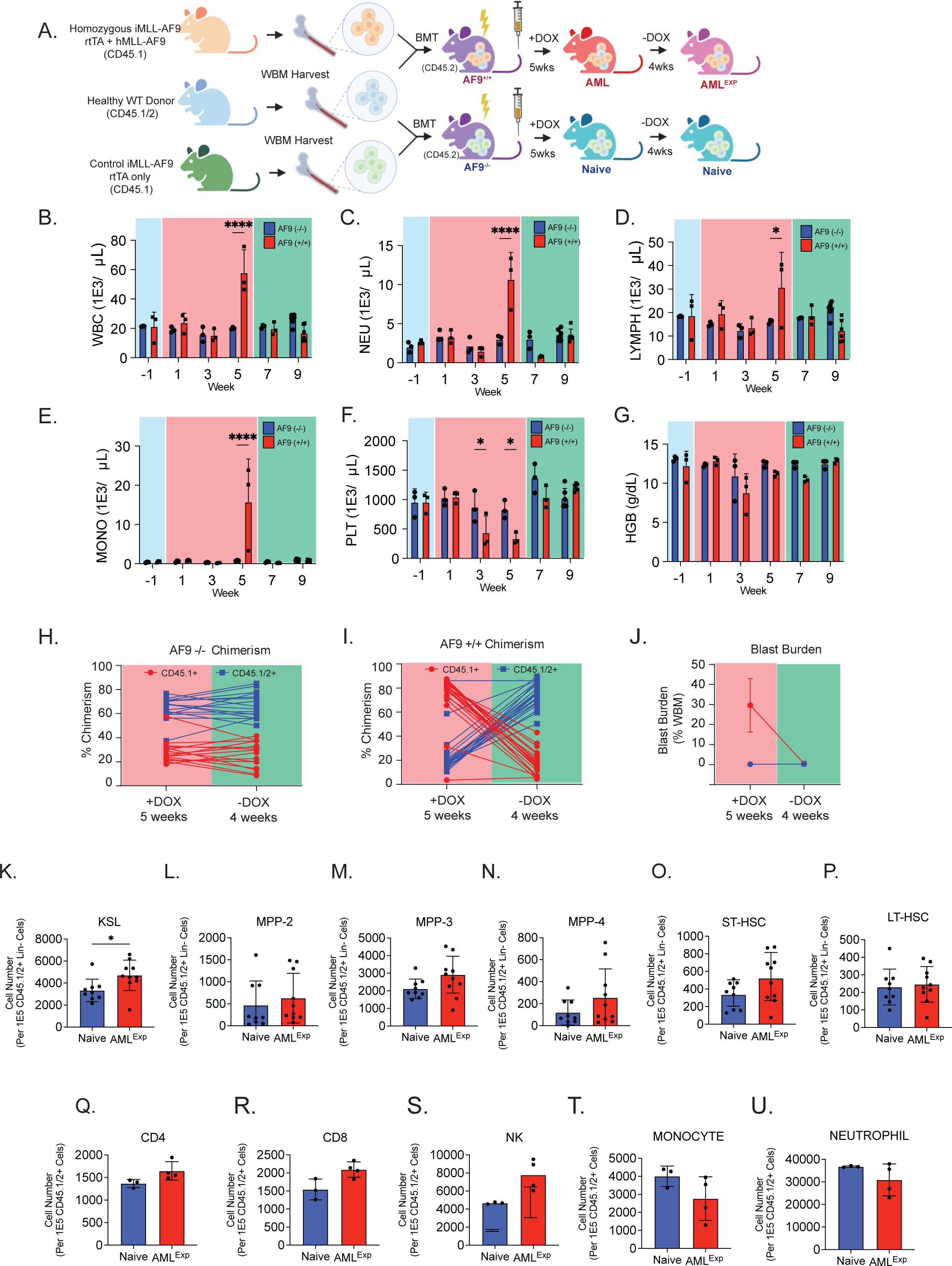
Generation of an AML remission model *in vivo.* (A) Schematic of the AML remission model generation and induction timeline. AF9^+/+^ mice was generated by co-transplanting WT healthy BM (CD45.1/2+) and non-induced homozygotes hMLL-AF9 BM (CD45.1+) into myeloablated recipients (CD45.2+). AML progression was initiated by subjecting mice to DOX water feed (1g/L) for 5 weeks. AML regression was achieved by withdrawing DOX water for 4 weeks. CBC was performed on AF9^+/+^ and AF9^-/-^ mice over the course of pre-induction (week -1; blue), AML progression (week 1, 3, 5; red), and remission (week 7, 9; green) (n=≥3). CBC counts in white blood cell (B), lymphocyte (C), neutrophil (D), and monocyte (E), platelets (F) and hemoglobin (G). Assessment of BM chimerism (H-I) and NGFR-hMLL-AF9-expressing cells (J) in AF9^-/-^ (H) and AF9^+/+^ (I) mice at 5 weeks +DOX and 4 week off-dox (n=≥3). SLAM compartment immunophenotyping was also characterized (K-P) including KSL (CD45.1/2+ Lin- cKit+ Sca1+), MPP-2 (CD45.1/2+ Lin- cKit+ Sca1+ Flk2- CD150+ CD48+), MPP-3 (CD45.1/2+ Lin- cKit+ Sca1+ Flk2- CD150- CD48+), MPP-4 (CD45.1/2+ Lin- cKit+ Sca1+ Flk2+ CD150- CD48+), ST-HSC (CD45.1/2+ Lin- cKit+ Sca1+ Flk2- CD150- CD48-), and LT-HSC (CD45.1/2+ Lin- cKit+ Sca1+ Flk2-CD150+ CD48-) (n=6). BM immune microenvironment population immunophenotyping (Q-U) was performed on common CD3 (CD45.1/2+ CD3+), CD4 (CD45.1/2+ CD3+ CD4+), CD8 (CD45.1/2+ CD3+ CD8+), NK ((CD45.1/2+ CD3- NK1.1+), common myeloid (CD45.1/2+ CD11b+), monocyte (CD45.1/2+ CD11b+ Ly6C+ Ly6G-), neutrophil (CD45.1/2+ CD11b+ Ly6G+ Ly6C-) (n=≥3). Values are expressed as mean ± standard deviation. Statistical significance was calculated using student’s t-test. *p<0.05; **p<0.01; ***p<0.001.

To track the progressive gain in leukemic burden upon DOX induction, we monitored complete blood counts (CBC) in chimeric mice, observing significantly elevated white blood cells in AF9^+/+^ mice compared with AF9^-/-^ controls (Figure 1B-E). A concomitant platelet count reduction was observed starting from week 3 and persists through week 5, whereas the reduction in hemoglobin levels was marginal, reflecting the longer half-life of red blood cells (Figure 1F-G). BM analysis of cohorts at week 5 revealed a significant medullary leukemic burden, measured by CD45 chimerism and NGFR^+^ frequency (Figure 1H-J). AML remission was achieved by withdrawing DOX and thereby extinguishing hMLL-AF9 oncogene expression, which leads blasts to differentiate^14^. BM analysis of AML^EXP^ mice 4 weeks off-DOX confirmed the complete absence of blasts, with WT/AF9 BM ratio and NGFR^+^ frequencies returning to baseline levels that mirror the AF9^-/-^ control (Figure 1H-J). Complete blood count analysis also showed a complete recovery of platelets and leucocytes to baseline, consistent with AF9^-/-^ control (Figure 1B-G).

To ascertain complete clearance, we investigated FACS-sorted AML blasts, from fully-induced homozygous hMLL-AF9 mice, and subjected the blasts to a DOX starvation challenge *ex vivo* (Supplementary Figure 1A). Placed in liquid culture, blasts in DOX-containing (+DOX) medium continue to proliferate, reaching 200-times their starting cell number over a 14-day period, while no proliferation and declining viability were observed in AML blasts cultured in control (-DOX) medium (Supplementary Figure 1B-C). Similarly, by expanding freshly harvested AML blasts in DOX- containing methylcellulose assay, we found significantly increased numbers of leukemia colony forming units (CFU-L) over 14 days, with no growth in the absence of DOX (Supplementary Figure 1D-G). Interval gene expression analysis of *ex vivo* blasts further confirmed that expression of the hMLL-AF9 fusion oncogene and its targets Meis1 and Hoxa9 becomes rapidly undetectable in the absence of DOX (Supplementary Figure 1H-M).

Given the dynamic changes in leukemic burden and associated inflammation, we assessed the immune cell composition in the BM niche of AML^EXP^ mice in remission. Results showed that common mature lymphoid CD4 (CD45.1/2+ CD3+ CD4+), CD8 (CD45.1/2+ CD3+ CD8+) NK (CD45.1/2+ CD3- NK1.1+) population (Figure 1K-N) and myeloid monocyte (CD45.1/2+ CD11b+ Ly6C+ Ly6G-), neutrophils (CD45.1/2+ CD11b+ Ly6G+ Ly6C-) populations (Figure 1O-Q) frequencies return to steady state baseline comparable to HSPC^naive^ controls. By contrast the AML^EXP^ BM revealed significantly elevated KSL immunophenotype (Lin- cKit- Sca1+) HSPC frequency compared to control, with a trend for MPP3 (myeloid-primed, inflammation prone), but otherwise balanced progenitor frequencies (Figure 1R-W). Concurrent multiplex secretome analysis, found that AML^EXP^ BM plasma did not exhibit consistent changes in a 44-plex panel of inflammatory chemokines or cytokines (Supplementary Figure 2A). Selective analysis of BM (Supplementary Figure 2B-G) and spleen (Supplementary Figure 2H-R) neutrophils and monocytes for interleukin (IL) -1β, IL-6 and TNFα confirms that gene expression returns to steady state control levels (Supplementary Figure 2B-G). Collectively, these findings demonstrate that the BM in AML^EXP^animals in remission returns to homeostatic state, resembling that of healthy naïve mice, without evidence for persistent inflammation in the BM and extramedullary. This state resembles the “resting period” of trained immunity.

### A distinctive transcriptomic signature and secondary inflammatory response in AML- experienced HSPC in remission

Given the initial sterile inflammatory response to AML blasts we reported in this model^12^ and the elevated KSL frequency among AML-experienced (HSPC^AML^) in remission seen here, we wanted to define the baseline transcriptome and then ask whether HSPC^AML^ would respond differently from HSPC^Naive^ to a secondary inflammatory challenge. To investigate, we subjected AML^EXP^ mice to a heterologous inflammatory challenge with LPS (AML^EXP^+LPS). LPS-challenged naïve mice served as control (Figure 2A). First, BM assessment of common mature lymphoid (Supplementary Figure 3A-D, mature myeloid (Supplementary Figure 3E-G), SLAM population (Supplementary Figure 3H- M), and BM plasma secretome (Supplementary Figure 3N) showed that immune cell composition and cytokine production in the BM of AML^EXP^ mice are phenotypically similar to Naïve mice. To further investigate whether distinct transcriptional differences are observable in HSPC^AML^, we performed an RNA-Seq analysis on FACS-purified HSPC from LPS-injected AML^EXP^ mice and naïve mice in remission. PBS-injected AML^EXP^ and Naïve mice served as controls. Animals were sacrificed, whole bone marrow enriched for KSL immunophenotype HSPC for RNA extraction and library construction to undergo bulk RNA-Seq. We observed global transcriptomic differences between the conditions, whereby HSPC^AML^ revealed 2,270 differentially expressed genes (DEGs), with 1214 upregulated and 1056 downregulated targets compared with PBS-injected HSPC^Naive^ (Figure 2B). This difference between experienced and naïve conditions became more pronounced following the LPS challenge, when both HSPC^AML^ and HSPC^Naive^ cohorts further adapt global gene expression (Figure 2C-D). This involved a greater number of genes in the AML experienced cohort with downregulated expression.

**Figure 2.**
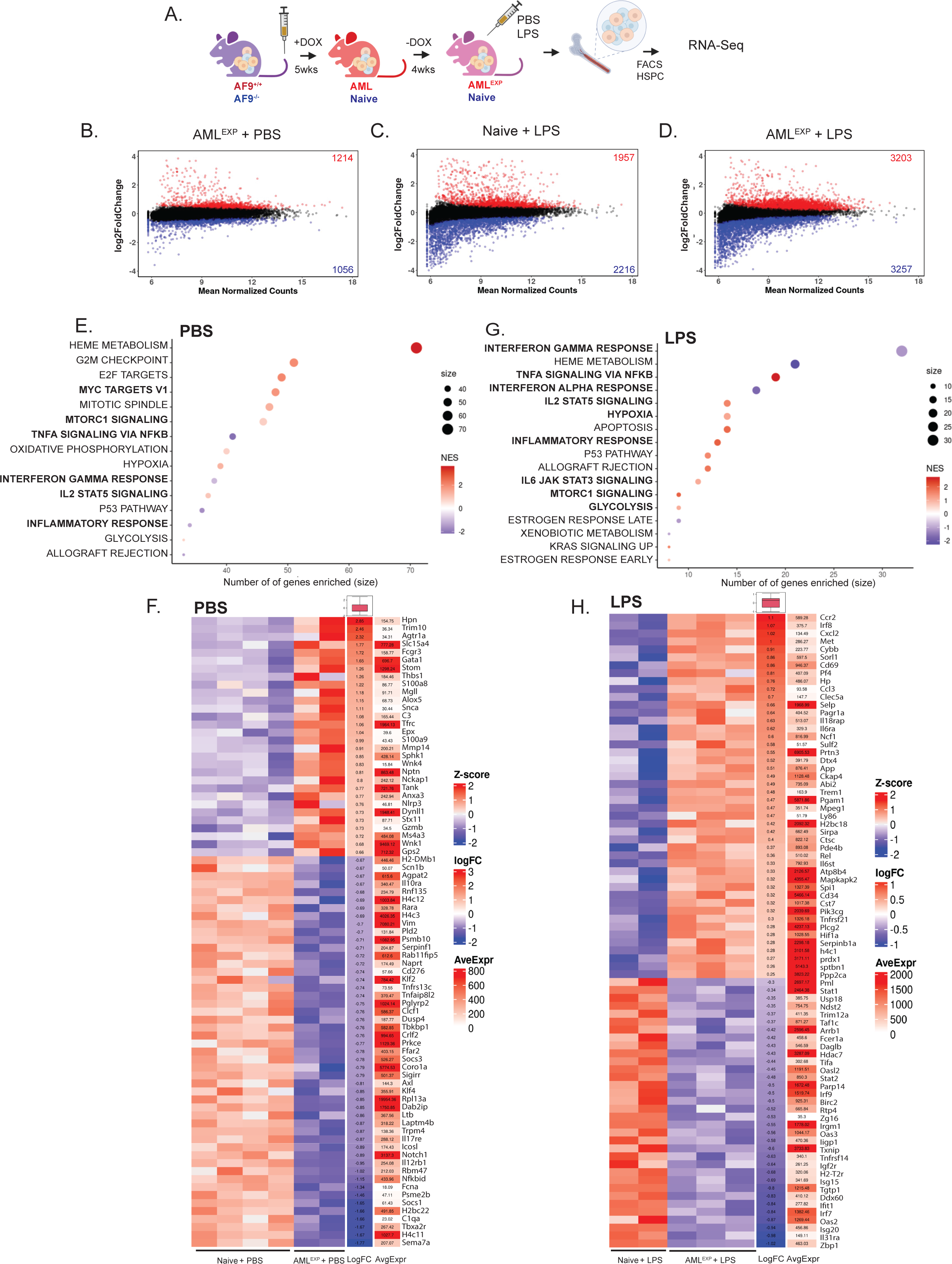
Distinct transcriptional signature in HSPC^AML^. (A) Schematic of experimental approach in assessing HSPC transcriptome at baseline (HSPC^AML+PBS^) or following LPS challenge (HSPC^AML+LPS^) in AF9^-/-^ (Naïve) and AF9^+/+^ (AF9-experienced; AML^EXP^) mice after 4 weeks into remission. FACS-sorted KSL (CD45.1/2+ Lin- cKit+ Sca1+) were subjected to RNA-Seq 24 hours after injection (n=≥3). (B) A global transcriptomic comparison was performed by comparing global differential gene expression analysis on AML^EXP^-HSPC (HSPC^AML^) injected with PBS (HSPC^AML+PBS^), HSPC^AML^ injected with LPS (HSPC^AML+LPS^), and Naïve HSPC injected with LPS (HSPC^Naive+LPS^) normalized to control Naïve HSPC injected with PBS (HSPC^Naive+PBS^). Further analysis of HSPC^AML+PBS^ against HSPC^Naive+PBS^ via gene set enrichment analysis (GSEA) showed several differentially enriched hallmark pathways (E) in HSPC^AML^ at baseline remission state. Differentially expressed inflammation-related genes were also identified (F). Analysis of HSPC^AML+LPS^ against HSPC^Naive+LPS^ also revealed differentially enriched pathways (G) and inflammation-related genes (H).

Next, to understand the biology reflected in these transcriptional differences in AML experienced HSPC following remission, we undertook gene set enrichment analysis (GSEA) and found HSPC^AML^ positively enriched for a number of metabolism relevant pathways including *hypoxia, heme metabolism, MYC* and *mTOR signaling*, a recurring pathway in other models of innate immune training (Figure 2E)^5,15,16^. Negatively enriched were key inflammatory pathways: *TNFα, Interferon-ψ* and overall *inflammatory response*. This was broadly consistent with the generally balanced BM secretome data and select transcriptional responses for IL-1β, TNFα and IL-6 in mature myeloid cells (Supplementary Figure 2A-G). Closer inspection of individual DEGs further identified several significantly genes in HSPC^AML^ that were upregulated at baseline, including members of the S100 family (*S100a8, S100a9*) and *Nlrp3*, all associated with inflammation, as well as *GATA1* encoding a core hematopoietic transcription factor. Also notable is a larger group of downregulated genes, including members of the SOCS family of cytokine signaling suppressors (*SOCS1, SOCS3*), but the transcription factor Klf4^17^ known to be downregulated in LPS sepsis and inflammation (Figure 2F).

When we interrogated the pathways enriched in HSPC^Naive^ and HSPC^AML^ following a LPS challenge (Figure 2G), we again observed positive enrichment in HSPC^AML^ related to inflammation (*TNFa signaling via NFkB, IL2-STAT5 signaling, inflammatory response, IL6-JAK-STAT3 signaling)* and the *mTORc1 signaling* and *glycolysis* pathways. By comparison, the Interferon Type 1 and -2 pathways (*Interferon alpha response* and *Interferon gamma response*) are negatively enriched in LPS-challenged HSPC^AML^, suggesting that AML experienced HSPC in remission respond selectively to a secondary heterotypic inflammatory stimulus. At the gene level, 46 inflammation related genes were significantly upregulated in HSPC^AML^ after LPS, prominently including *Il6ra, Met* and *Cxcl12, Ccl3, Pf4 and SelP*, that are all extensively annotated for their role in inflammation (Figure 2H). Our finding demonstrates that KSL-specified HSPC not only respond differently to sterile inflammatory exposure in the AML niche, but their gene expression is further biased toward inflammation after a heterologous secondary stimulus. The constitutive upregulation of general inflammatory pathways and specific stimulation of TNFα (*TNFα signaling via NFkB* hallmark pathway) in HSPC^AML^ indicates a heightened secondary inflammatory response.

### An altered chromatin accessibility landscape in HSPC^AML^ persists in remission

Altered chromatin accessibility and metabolic activation are hallmarks of the HSPC memory in other models of trained immunity where chromatin accessibility enables rapid and heightened responses to secondary stimuli. To ask whether the systematic changes in inflammatory and metabolic gene expression programs in HSPC after exposure in the AML niche is matched by an altered chromatin landscape in remission, we performed an assay for transposase-accessible chromatin with sequencing (ATAC-Seq) on DNA from KSL sorted HSPCs of animals in remission (Figure 3A). Following generation of AF9^(+/+)^ hybrid animals, DOX induction and monthlong remission, the AML experienced animals were sacrificed 24 hours after receiving PBS or LPS injections, for marrow harvest, HSPC enrichment (KSL immunophenotype) and construction of libraries. Through differential accessibility (DA) analysis of the ATAC-Seq data across promoter and putative enhancer regions (Figure 3B), we identified 283 differentially accessible peaks in HSPC^AML^ at baseline compared to HSPC^Naive^ controls, with 172 peaks that are more accessible (Figure 3C). A pathway analysis of DAs shows negative, non-significant, enrichment for pathways relating to inflammation (*TNFa signaling via NFkB, IL2-STAT5 signaling, interferon gamma response, inflammatory response, mTORc1 signaling* and *glycolysis)* (Figure 3D).

**Figure 3.**
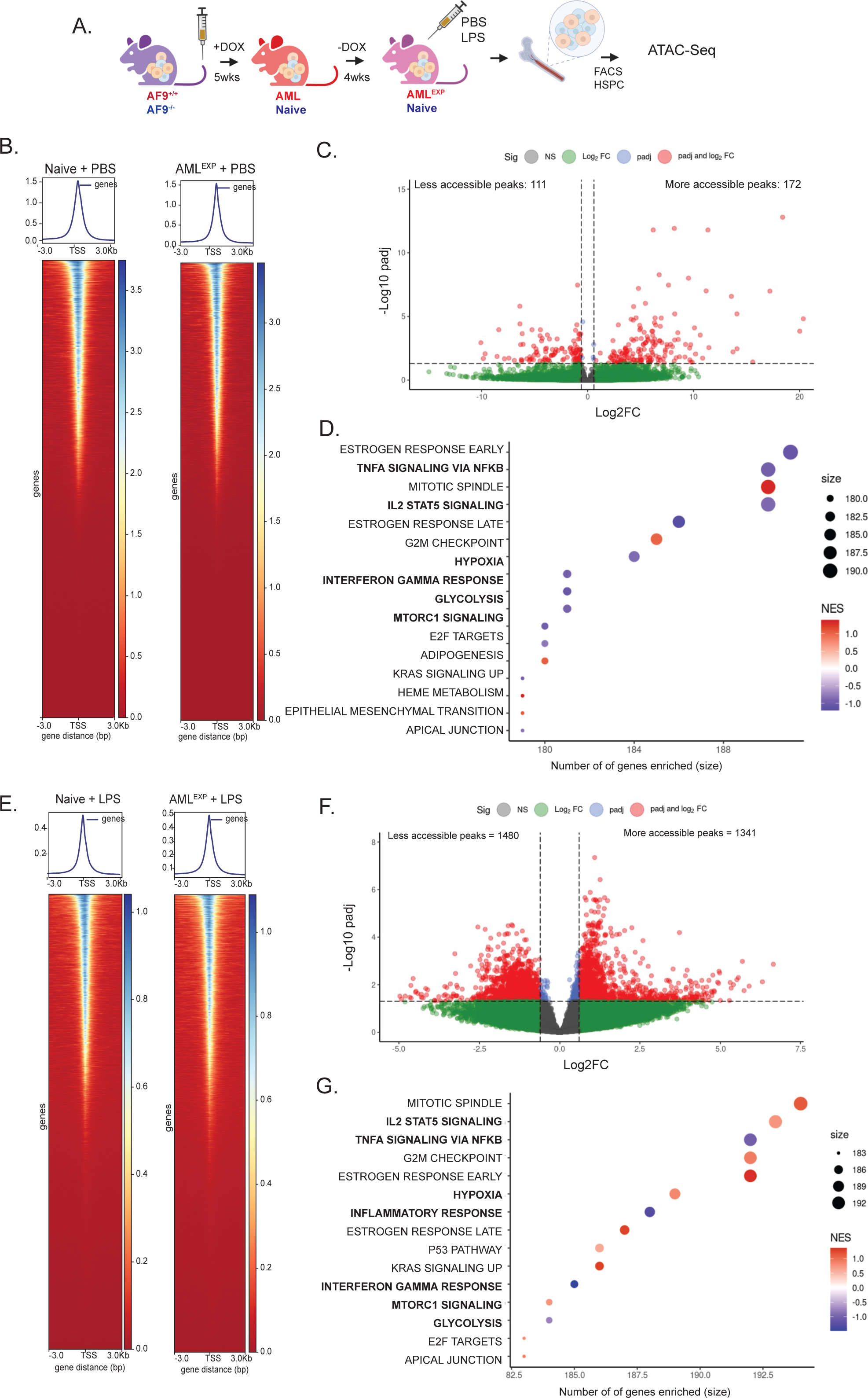
Distinct chromatin accessibility signature in HSPC^AML^. A) Schematic of experimental approach in assessing HSPC chromatin accessibility at baseline (+PBS) or following LPS challenge (+LPS) in Naïve and AML^EXP^ mice after 4 weeks into remission. FACS-sorted KSL (CD45.1/2+ Lin-cKit+ Sca1+) were subjected to ATAC-Seq (n = ≥3) 24 hours after injection. (B) To assess chromatin accessibility in HSPC^AML^ at baseline AML remission, differentially accessible peaks were called in HSPC^AML+PBS^ and HSPC^Naive+PBS^ and visualized (C). GSEA analysis of the genes associated with the differentially accessible peaks revealed differentially expressed pathways relating to inflammation and metabolism (D). Chromatin accessibility was also assessed in LPS-challenged HSPC^AML^ (E) with differentially accessible peaks identified (F) and pathways enriched (G).

To test whether a secondary challenge would alter chromatin accessibility in HSPC^AML^, animals in remission that received a single dose of LPS were compared to naive control animals (HSPC^AML^ and HSPC^Naive^) to ATAC-Seq analysis (Figure 3E). By contrast to PBS controls, we observed more substantial changes in accessibility, particularly toward increased significance of accessibility changes and a substantially increased number of DA peaks (n=2831) in LPS-challenged HSPC^AML^ compared to control (Figure 3F). Through GSEA analysis, we found pathways associated with inflammation (*IL2-STAT5 signaling, TNFa signaling via NFkB, inflammatory response, and interferon gamma response),* along with *mTOR*c1 signaling and *glycolysis* that are most differentially enriched compared with baseline HSPC^AML^ (Figure 3G). Several pathways stand out for their relationship with inflammation in HSPC. For example, WT KRAS signaling reinforces inflammation and promotes recovery of HSPC function after myelosuppression^18^. The prominent estrogen response (including both early and late pathways) is particularly striking, likely reflecting the increased sensitivity in female derived grafts, but especially consistent with the role of estrogens in regulating innate immune cells^19^. Incidentally, chromatin accessibility associated with the G_2_M checkpoint sites was previously identified in the ATAC-Seq dataset from a study of LPS mediated innate immune training^20^. Together, AML-mediated chromatin modifications in HSPC^AML^ are evident in remission and become more prominent after a heterologous LPS inflammatory challenge.

### Correlative analysis of the inflammatory response in AML experienced HSPCs

To better understand the overall dynamics underlying inflammatory reprogramming and recall responses by LPS-challenged HSPC^AML^, we performed a correlative RNA/ATAC-Seq analysis to identify common active DGEs and DARs. At baseline, a total of 674 genes scored as both differentially expressed and differentially accessible (DGE+DAR) in HSPC^AML^ at baseline with *heme metabolism* and *fatty acid metabolism* positively enriched (among 372 sites) and *inflammatory response*, *TNFα via NFκB* and *IL-6* signaling among 302 negatively enriched sites (Supplementary Figure 4A-B). When remission animals were challenged with LPS, the expression of 372 genes was upregulated with more accessible chromatin at the corresponding promoter/enhancer sites, while 355 genes showed concomitant loss of expression and less accessible chromatin (Figure 4A). Among the latter, there was a specific loss of expression and accessibility in *Ifn-α response* and *Ifn-ψ response* pathways, and significant consensus gains in *mTORC1* as well as *inflammatory response* pathways, further including *IL-6-JAK-STAT signaling* and *TGF-b signaling* (Figure 4B). At the level of individual genes, the following inflammation-related genes are upregulated in HSPC^AML^ after LPS challenge (*Ccr2, Irf8, Cxcl2, Met, Cybb, Sorl1, Cdkn1a, Cd69, Pf4, Hp, Ccl3, Clec5a, Selp, Flt3, Pagr1a, Il18rap, Il6ra, Ncf1, Sting1*). Among these, six genes (*Ccr2, Irf8, Met, Sorl1, Pf4, and Hp*) are also associated with more accessible chromatin (Figure 4C). Notably, several glycolysis related genes (*Met, Egln3, Pgam1, Pkm, Sdhc, Pgk1, Pgk1, Ier3, and Hif1a)* were significantly upregulated (Supplementary Figure 5A).

**Figure 4.**
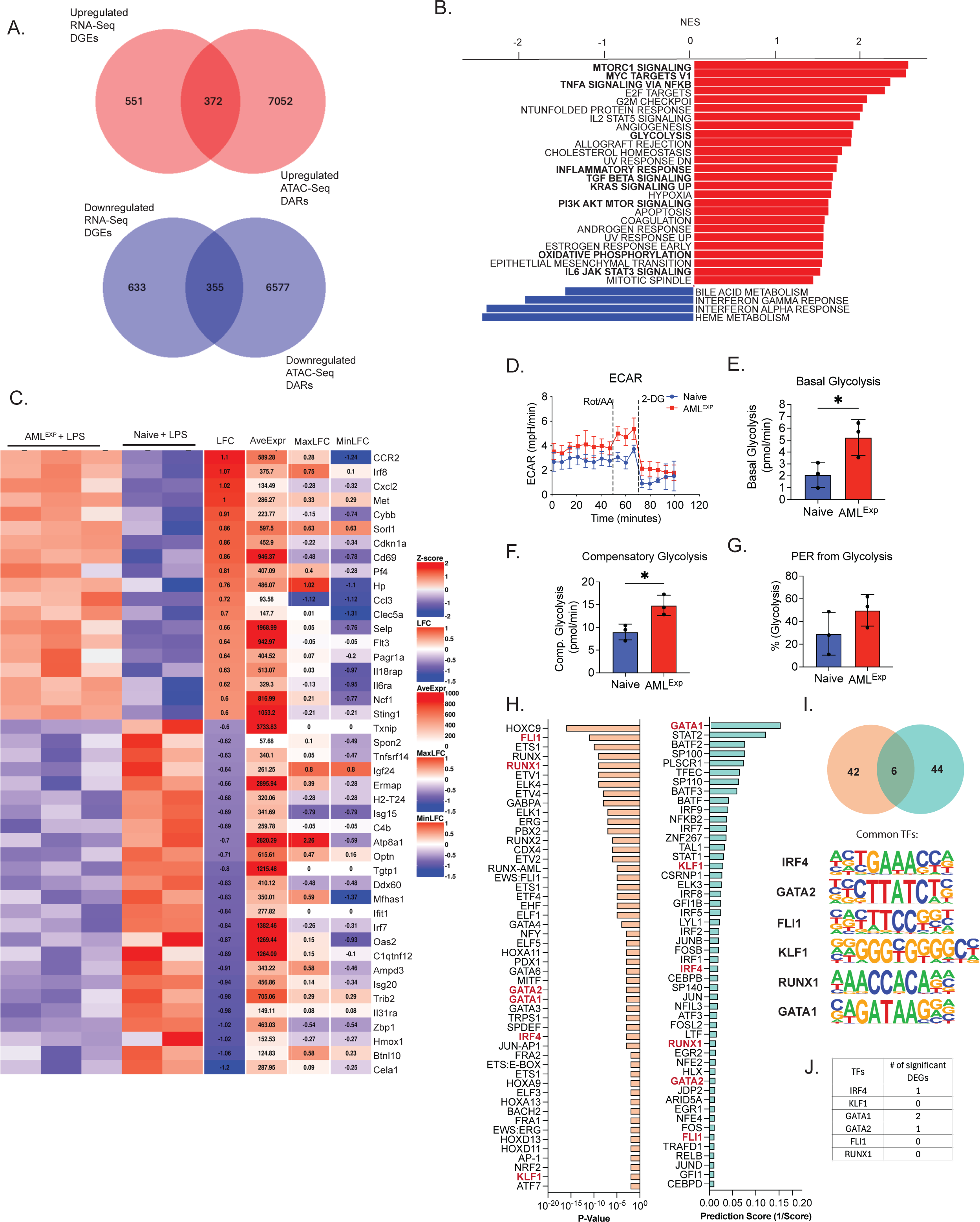
Correlative RNA-Seq/ATAC-Seq analysis of HSPC^AML^ challenged with LPS. (A) Transcriptomic and chromatin accessibility correlative analysis between HSPC^AML+LPS^ and HSPC^Naive+LPS^ revealed 727 genes that are both differentially expressed and accessible. (B) GSEA analysis of these genes resulted showed significant number of pathways that are positively enriched in HSPC^AML+LPS^. (C) Top differentially expressed inflammation-associated genes (LFC, RNA-Seq) and their corresponding maximum (MaxLFC) and minimum (MinLFC) LogFC in ATAC-Seq. (D) Metabolic assessment of FACS-purified HSPC^AML^ was evaluated using Metabolic Rate Assay (Seahorse) (n=3), and its corresponding measurements in (E) basal glycolysis, (F) compensatory glycolysis, and (G) the percentage of proton efflux rate (PER) from glycolysis. Comparison of top 50 motifs predicted in HSPC^AML+PBS^ based on chromatin accessibility (H, left, ranked based on adjusted p-value from low to high) and top 50 transcription factors (TFs) predicted based on HSPC^AML+LPS^ gene expression (H, right, ranked based on prediction score from high to low) led to identification of 6 TFs (I) that are commonly annotated. (J) Number of known target genes associated with each of the TFs that are differentially expressed in HSPC^AML+LPS^. Values are expressed as mean ± standard deviation. Statistical significance was calculated using student’s t- test. *p<0.05.

HSC quiescence relies on aerobic glycolysis^21^. A recurring theme in our dataset investigating AML- experienced HSPC in remission, but especially in the correlative analysis, was the enrichment for metabolic activity, including *glycolysis*, *oxidative phosphorylation*, *hypoxia, PI3K-mTORc1* and *mTORc1 signaling* pathways (Figure 4B). To functionally validate the metabolic state of HSPC in AML remission, we next undertook real-time metabolic analysis using the seahorse assay. Using FACS-sorted KSL cells (4x10^4^ cells per well) from animals in remission (no LPS challenge) we performed canonical study sequences for metabolic flux and oxygen consumption. Here, we found that basal and compensatory glycolytic rates are significantly higher in HSPC^AML^ compared to HSPC^Naive^ (Figure 4C-F). Several glycolysis related genes were significantly upregulated (*Hmmr, Gpc4, Gnpda1, Hk1*) in HSPC^AML^ (Supplementary Figure 5B). When we tested for mitochondrial function in HSPC^AML^ using the Mito Stress assay, we observed no significant difference in oxidative phosphorylation in HSPC^AML^ (Supplementary Figure 5C-E). Consistent with our observation of gains in chromatin accessibility at genes involved in glycolysis, the data reveal that altered metabolic activity coincides with epigenetic modification in HSPC^AML^. Real-time metabolic activity measurement in HSPC^AML^ after LPS stimulation did not detect additional shifts in glycolytic (Supplementary Figure 4F-I) or oxidative phosphorylation activities (Supplementary Figure 5J-L).

Hematopoiesis regulation relies heavily on a core set of transcription factors (TF) and here we probed for the potential involvement in the inflammatory reprogramming of AML experienced HSPCs. To investigate this, we first performed Hypergeometric Optimization of Motif EnRichment (HOMER) DNA binding motif discovery analysis in the ATAC-Seq dataset to identify open chromatin regions present in HSPC^AML^ at baseline. Reasoning that open TF target sites in remission animals are more likely to express during a subsequent challenge, we probed the RNA-Seq data from LPS- challenged HSPC for reference by subjecting significantly DEGs in HSPC^AML^ to ChIP-X Enrichment Analysis 3 (ChEA3) analysis, a gene expression-based TF enrichment prediction tool^22^. Comparing the top 50 motif binding sites predicted by HOMER in HSPC^AML^ at baseline with the top 50 TF predicted in LPS-challenged HSPC^AML^ (Figure 4H), we identified 6 candidate TFs (IRF4, GATA2, FLI1, KLF1, RUNX1, GATA1) most likely to be involved (Figure 4I). To further identify which TFs are most involved, we queried the gene expression levels of putative targets associated with each TF. Among these, the GATA1 targets *Epb42*, *Spi1*, and *Pf4* are the most upregulated specific genes in HSPC^AML^ after LPS challenge (Figure 4J), making GATA1 a putative TF involved in driving leukemic inflammatory responses in trained HSPC^AML^.

## Discussion

Work over the past decade reveals that progenitor and mature myeloid cells can develop a memory of infection or experimental sterile inflammatory stimulation that augments their response to heterologous subsequent challenges^2^. Yet, unlike genetic re-combination by which adaptive immune cells selectively respond, retain and propagate a memory of prior exposure, innate immune training that confers heightened responsiveness to subsequent inflammatory challenges involves an epigenetic mechanism. The translational impact of these observations is only beginning to emerge, and here we demonstrate for the first time that the sterile inflammation that is a constitutive aspect of cancer niche similarly endows HSPCs with an adapted (i.e. trained) status.

Sterile fetal inflammation is required for physiologic hematopoietic specification *in utero* and serves as a self-limiting protective mechanism throughout life^23^. However, chronic inflammation initiates a process that recruits HSPCs, leading to an aging-associated myeloid bias in the compartment, clonal hematopoiesis and potentially HSC pool exhaustion^24^. In AML, inflammation is a constitutive feature of leukemogenesis that promotes disease progression and suppresses healthy hematopoiesis^9,25^. Little is known about the long-term impact on HSPC fate and function, as AML patients return to steady state hematopoiesis when they enter remission. Clinical observations in AML patients have revealed a significantly increased prevalence of CHIP, where clonal populations with enhanced fitness selectively expand in a self-reinforcing loop of chronic inflammation, that leads to systemic co-morbidities and a heightened risk for progression to MDS or AML^26,27^. Accordingly, observations of inflammatory reprogramming of long lived progenitors after mycobacterial infection, BCG- vaccination or β-glucan treatment have generated considerable interest in exploring cancer as a possible pathophysiologic source of HSPC memory and innate training.

We recently showed that HSPCs actively contribute to the inflammatory secretome in a translationally relevant murine model of AML^12^, but durable consequences for inflammatory recruitment of HSPCs in the cancer niche after disease remission have not been investigated. Here, we describe a novel chimeric model that enables timed expansion and ablation of the leukemic population without disrupting the shared BM niche or the added experimental variable of treatment- related inflammation. We show that these hematopoietic chimera -like AML patients- develop peripheral blood cytopenias during AML progression that resolve in remission (i.e. off DOX). Like a recent study using β-glucan to confer cTI, we find that both peripheral blood myeloid and lymphoid populations as well as BM progenitor distribution return to their steady state baseline, except for a persisting increase in KSL progenitors^28^. The remission status in the chimera also reveals the resolution of significant initial inflammatory signaling we reported in this model at diagnosis^12^, with modest, transcriptional evidence of inflammation. This is reminiscent of other non-cancer models of innate immune training^20,29^. Indeed, average DGE for most targets is less than 2-fold, and highly reproducible, with Ifn-ψ and Tnf-α GEO Hallmark inflammatory pathways both negatively enriched. This validates our earlier results in a cell line model of AML, where we tracked AML-experienced HSPC for 16 weeks after adoptive transfer to a naïve BM niche and showed a negative enrichment of inflammatory response pathways, but highly significant transcriptional activation across multiple core metabolic pathways, including *MYC-*, *mTORC1-*, *fatty acid metabolism*^30^.

The current, more translationally oriented, chimeric model also confirms a persisting metabolic signature evident in healthy HSPC in disease remission. Those transcriptomic results of a persistent metabolic phenotype in remission are reinforced by metabolic flux measures that reveal increased glycolysis using seahorse analysis and the correlative analysis of transcriptional and ATAC Seq data in this study. The data point to an involvement of mTOR signaling in AML training of HSPC, thereby echoing other non-cancer models of cTI ^5,6,16^. A similar concurrent shift toward glycolysis has been seen in trained monocytes^15^. In this regard the inferred hematopoietic transcription factors from our correlative analysis of transcriptome and accessible chromatin are notable for their consistent role in regulating metabolism. For example, GATA-, RUNX- and KLF1 TFs all contribute to metabolic shifts to glycolysis in mature and progenitor hematopoietic cells. Likewise, MYC and IRF4 link to metabolic reprogramming of myeloid cells ^31^. Perhaps most robustly, FLI1 has well annotated roles in regulating glycolysis in healthy HSPCs^32^.

The translational impact of our study is strongly supported by a recent study of AML patients (with cohort validation in additional datasets) that identified inflammation as a highly significant risk factor for a poor patient prognosis^11^. More studies will be critical to dissect which of patients sustain a chronic inflammatory environment beyond remission and how this shapes mortality in terms of relapse, drug resistance and CHIP evolution. Related, another intriguing recent study identifies a novel inflammation biased HSC subtype where a population of functionally unique pro-inflammatory HSC memory through epigenetic reprogramming^33^ and that aging associated myeloid biased phenotype is defined by unique transcriptome and function ^33,34^.

Adaptive immunity provides powerful protection from infections and has been harnessed for the treatment of cancer. Until recently, the evolutionarily older innate immune system was widely seen as more static. Altogether, our results suggest that AML experienced HSPCs acquire several hallmark features of the adapted (trained) immune state. More broadly, the US Centers for Disease Control and Prevention report nearly 2.0 million cases of cancer in 2024 ^35^. Here, we show for the first time that leukemia associated inflammation can durably reprogram healthy HSPC. We propose that cTI represents a poorly understood cause of morbidity and an untapped therapeutic approach to prevent or treat recurrent cancer.

### Limitations of our study

The objective in this study was to demonstrate that cancer associated sterile inflammation can reprogram HSPCs. While representative and widely validated our choice of a murine model of AML as an experimental model represents a specific AML subtype and results cannot be extrapolated to cancer without validation. Like most studies in the field, we focus on murine models and acknowledge the need to evaluate innate HSPC training in patient derived cells. The cTI paradigm posits that different secondary -and typically heterologous- stimuli can lead to different outcomes, that are alternatively pro-inflammatory or tolerizing. Consistent with many other studies, we have only evaluated LPS as a secondary challenge. Future studies will evaluate other inflammatory stimuli to determine the ability to rationally direct responses for therapeutic purposes. Finally, our work over the past decade characterized the role of EVs in the leukemic marrow crosstalk. Here, we did not explore individual EV components for their specific contribution to cTI.

## Acknowledgement

We gratefully acknowledge funding support from Alex’s Lemonade Stand Foundation for Childhood Research (DWC; Young Investigator Grant; 21-23996), Cure4Cam Childhood Cancer Foundation and Marshall County Childhood Cancer Awareness Corporation. We are indebted to Dr. Wei Tong and Nikolai Vantsev for access and guidance in using the Seahorse instrument.

## Data Availability

RNA-Seq and ATAC-Seq data will be deposited and become publicly available in the Gene Expression Omnibus (GEO) Database at the time of publication.

## Methods

### Animal Husbandry

Inducible (i)hMLL-AF9 mice were generously gifted by Dr. Shangqin Guo (Yale University). Chimeric AF9+/+ mice were generated by co-transplanting whole bone marrow from non-induced homozygotes iMLL-AF9 mice (CD45.1; homozygotes for rtTA and hMLL-AF9; 2E5 cells per recipient) and WT CD45.1/2+ mice (8E5 cells per recipient) into lethally irradiated C57BL/6J mice (CD45.2). Control AF9^-/-^ mice served as control and was generated by co-transplanting whole bone marrow from hMLL-AF9-null (CD45.1; homozygotes for rtTA only) and WT CD45.1/2+ mice. To induce leukemogenesis, chimeric mice were subjected to doxycycline (DOX; 1g/L supplemented with sucrose 10g/L) water feed for 5 weeks. To model experimental remission, DOX water feed was removed and replaced with normal water for 4 weeks. Animal experiments were approved and conducted in accordance with the Children’s Hospital of Philadelphia Institutional Animal Care and User Committee.

### Hematopoietic stem and progenitor cell harvest and isolation

Upon sacrifice, mouse bone marrow cells were collected from either flushing the tibias and femurs from hind legs alone, or in addition to crushing sternum and spine. Cells were collected in Hank’s Balanced Salt Solution (HBSS) supplemented with 1% P/S and filtered through cell strainer to generate single-cell suspension. Lineage negative cells were enriched using EasySep Mouse Hematopoietic Progenitor Cell Isolation Kit (Stem Cell Technologies), followed by staining with appropriate antibodies. Cells were sorted using FACSAria Fusion sorter (Becton Dickinson, Ranklin Lakes, NJ, USA).

### Methylcellulose colony cultures

AF9 blasts were FACS-sorted (NGFR+ cKit+ Gr-1+ Mac1+) from BM of fully-induced homozygotes hMLL-AF9 mice at moribund. FACS-sorted blasts were cultured *ex vivo* in StemSpan medium supplemented with Tpo (50ng/mL; Peprotech), Flt3 Ligand (50ng/mL; Peprotech), Scf (100ng/mL; Peprotech), and doxycycline (2μg/mL). Methylcellulose colony forming unit (CFU) assay were performed on AF9 blasts using MethoCult M3434 (Stem Cell Technologies) supplemented with (or without) doxycycline (2μg/mL) at a seeding density of 1E5 cells per well in 6-well plate. All cells were cultured in humidified incubator at 37 degrees Celsius and 5% CO_2_. CFU assays were imaged at day 7 and 14 using STEMVision (Stem Cell Technologies).

### Flow-cytometry

Flow cytometric analysis or cell sorting (FACS) were performed using either FACS Canto II or FACSAria Fusion Sorter (BD). Analysis of common mature lymphoid and myeloid in BM were performed on either whole BM flush or BM aspirates from hind legs. Cells were defined as follow: CD4 (CD3+ CD4+), CD8 (CD3+CD8+), NK (CD3-NK1.1+), Monocyte (CD11b+ Ly6C+ Ly6G-), Neutrophil (CD11b+ Ly6G+ Ly6C-). Samples for SLAM population analysis and cell sorting were first subjected to lineage-negative population enrichment using the EasySep Mouse Hematopoietic Progenitor Cell Isolation Kit (Stem Cell Technologies) according to manufacturer’s protocol. Cells were defined as follow: KSL (Lin- cKit+ Sca1+); MPP-2 (Lin- cKit+ Sca1+ Flk2- CD150+ CD48+); MPP-3 (Lin- cKit+ Sca1+ Flk2- CD150- CD48+); MPP-4 (Lin- cKit+ Sca1+ Flk2+ CD150- CD48+); ST-HSC (Lin- cKit+ Sca1+ CD150- CD48-); and LT-HSC (Lin- cKit+ Sca1+ CD150+ CD48-).

### RNA-Seq

RNA was isolated from HSPCs using RNeasy Plus Micro kit and quantified and analyzed using NanoDrop and TapeStation System. Library was prepared using Illumina Stranded Total RNA Prep Kit according to manufacturer’s protocol and sequenced using NovaSeq 6000. Sequence read preprocessing, including quality control, mapping, and expression quantification, was performed using the nf-core/rnaseq pipeline v3.12.0 (Ewels et al., 2020). Read quality was assessed using FASTQ v0.11.9 (https://www.bioinformatics.babraham.ac.uk/projects/fastqc/), and adaptors/low- quality bases were trimmed using Trim Galore v0.6.7 (https://www.bioinformatics.babraham.ac.uk/projects/trim_galore/). The resulting high-quality read fragments were mapped onto the GRCm38 mouse reference genome using the STAR aligner v2.7.10a (Dobin et al., 2013). Then, gene-level abundance from Ensembl comprehensive annotation was estimated using RSEM v1.3.1 (Li and Dewey, 2011). DESeq2 v1.38.3 (Love et al., 2014) was used to generate variance stabilized transformed (VST) normalized expression count matrices and to assess differential gene expression between AML-experienced and naive HPCSs with either PBS or LPS treatment. Transcription factor prediction analysis were performed utilizing the web-based ChIP-X Enrichment Analysis (ChEA3) tool (https://maayanlab.cloud/chea3/) ^22^. Using significantly differentially expressed genes as input, a list of predicted TFs were generated and ranked based on the prediction score calculated by the algorithm.

### ATAC-Seq

Genomic DNA was isolated from HSPCs, and the ATAC-Seq library was prepared according to OMNI-ATAC protocol as previously described, followed by 50 bp paired-end sequencing using Illumina NovaSeq 6000. Data preprocessing, including quality control, read mapping, and peak calling, was carried out using the nf-core/atacseq (v2.0). Quality checks were performed on raw sequence data using FASTQC, and adaptors and low-quality bases were removed using Trim Galore. Trimmed reads were then aligned to the mouse reference genome (GRCm38) using BWA (Li & Durbin, 2009) and filtered to remove PCR duplicate reads using Picard MarkDuplicates (v2.27.4) and mitochondrial DNA. Peaks were called using MACS3 (Zhang et al., 2008) after excluding the ENCODE blacklist, low-complexity regions in the mouse genome (Amemiya, Kundaje, & Boyle, 2019). Consensus peak sets consistent among biological replicates determined for each sample using the BEDTools (v2.30.0) intersect function (Quinlan & Hall, 2010), and the Subread featureCounts v2.0.1 (Liao et al., 2014) used to quantify consensus peak intervals read counts. Consensus peaks were annotated with the ChIPSeeker v1.40.0 (Yu et al., 2015) using the latest Ensembl mouse reference gene set to define promoter regions around transcription start sites. Differentially accessible regions (DARs) between AML-experienced and naive HPCSs with either PBS or LPS treatment were determined using limma-voom v3.60.2 (Ritchie et al., 2015) following sample batch correction on the raw consensus count matrix using ComBat-Seq (Zhang et al., 2020) and normalization with edgeR calcNormFactors v4.2.0 (Robinson & Oshlack, 2010) on log2 counts per million reads (CPM). Enrichment of motifs associated with DARs (adj.P.Val < 0.05) was performed using HOMER (Heinz et al., 2010).

### Gene Set Enrichment

Gene set enrichment analysis (GSEA) was performed using fgsea (v1.30.0) for with differential gene expression and differential accessibility analyses results on the Molecular Signatures Database (MSigDB) mouse collections for Hallmark, Reactome, and Gene Ontology functional categories, as well as a subset pathways genes associated with inflammatory and immune response, metabolism, chromatin and epigenetic, and signaling pathways to identify enriched pathways (Liberzon et al., 2015).

### Seahorse Assay

Real-time metabolic activity of HSPCs were measured using Seahorse XF HS Mini Analyzer (Agilent). Briefly, 50,000 HSPCs were seeded into each well of the XF HS PDL Miniplates. Glycolytic Rate and Mito Stress assays were performed according to manufacturer’s protocol.

## Data Availability

RNA-Seq and ATAC-Seq data will be deposited and become publicly available in the Gene Expression Omnibus (GEO) Database at (https://www.ncbi.nlm.nih.gov/geo/).

---

**Supplementary Figure 1.**
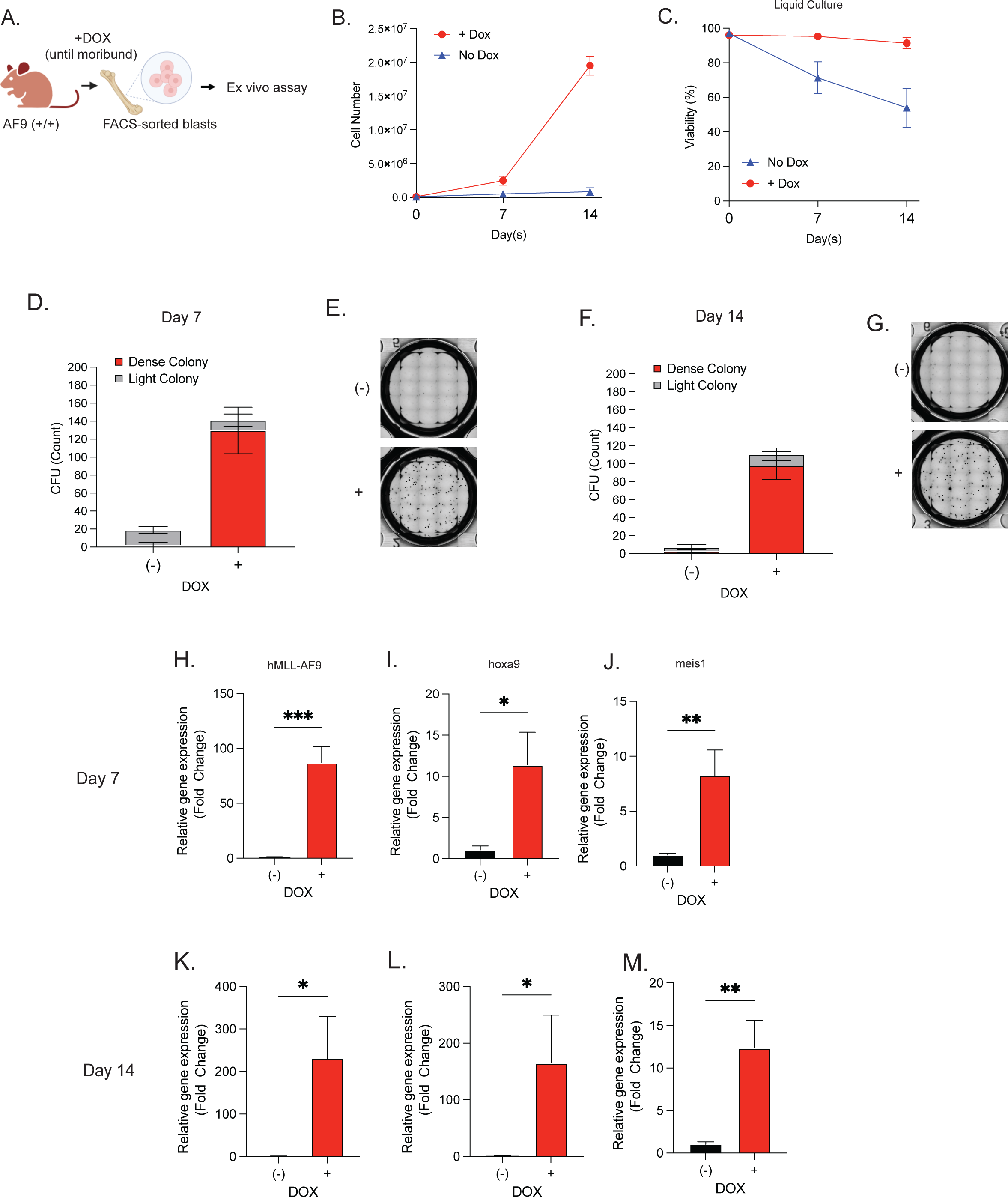
Characterization of AML blast fate with doxycycline starvation challenge. (A) AML blasts were FACS-sorted from the BM of fully induced AML at moribund, and blasts were cultured *ex vivo* in liquid culture (with/without DOX). Cell number (B) and viability (C) of blasts were tracked over the course of 14 days (n=3). AML blasts were also subjected to colony forming unit assay (with/without DOX) and colony numbers were visualized and counted at day 7 (D-E) and day 14 (F-G) (n=3). Oncogene expression analysis of AML blasts at day 7 (H-J) and day 14 (K-M) *ex vivo* (n=3).

**Supplementary Figure 2.**
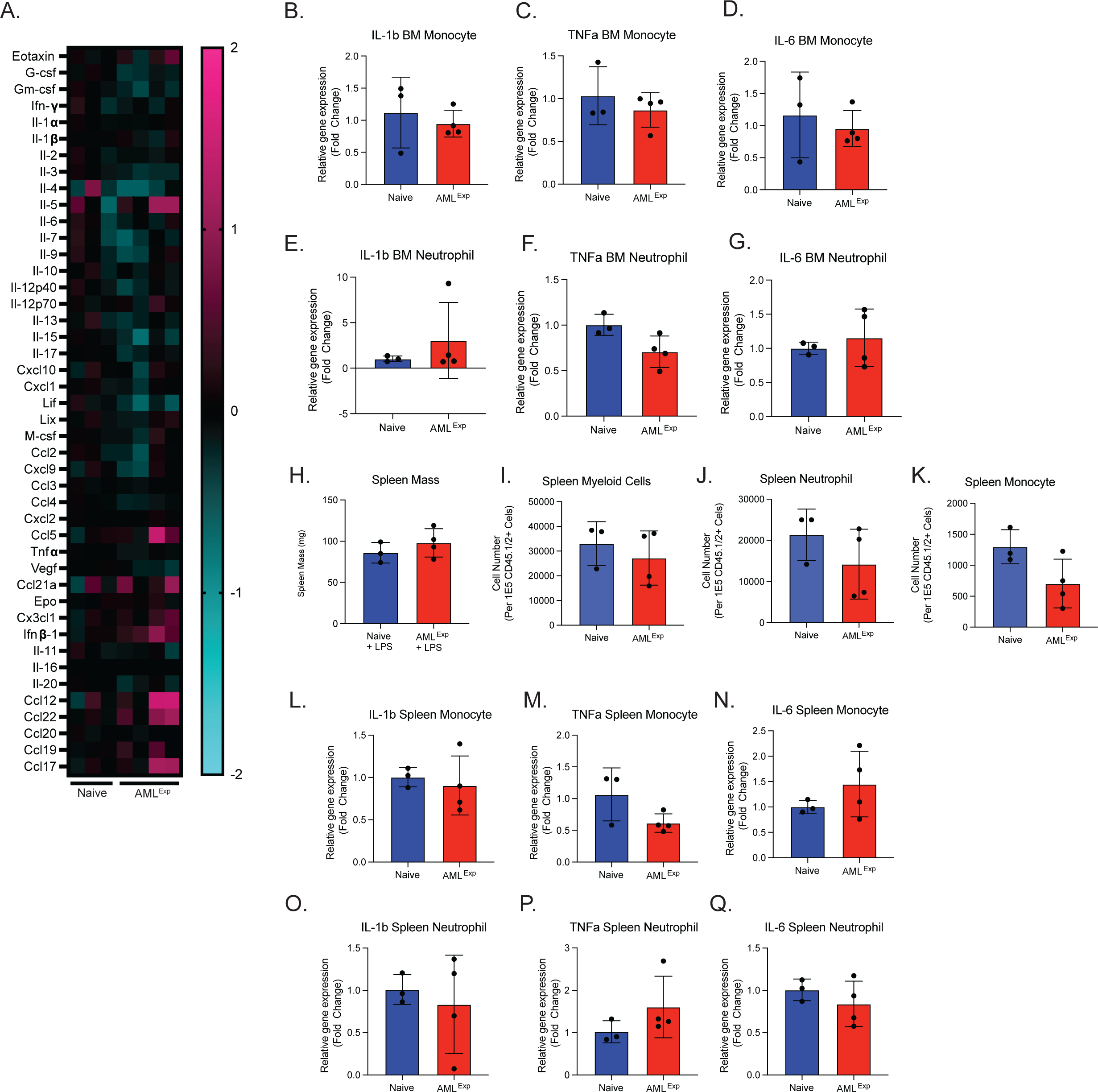
Cytokine profiling of BM plasma in AML^EXP^ mice. (A) 44-Plex chemokine/cytokine discovery assay analysis of BM plasma from AML^EXP^ and Naïve mice (n=≥3). Gene expression of select cytokine-encoding transcripts in BM monocyte (B-D) and neutrophils (E- G) (n=3). Statistical significance was calculated using student’s t-test.

**Supplementary Figure 3.**
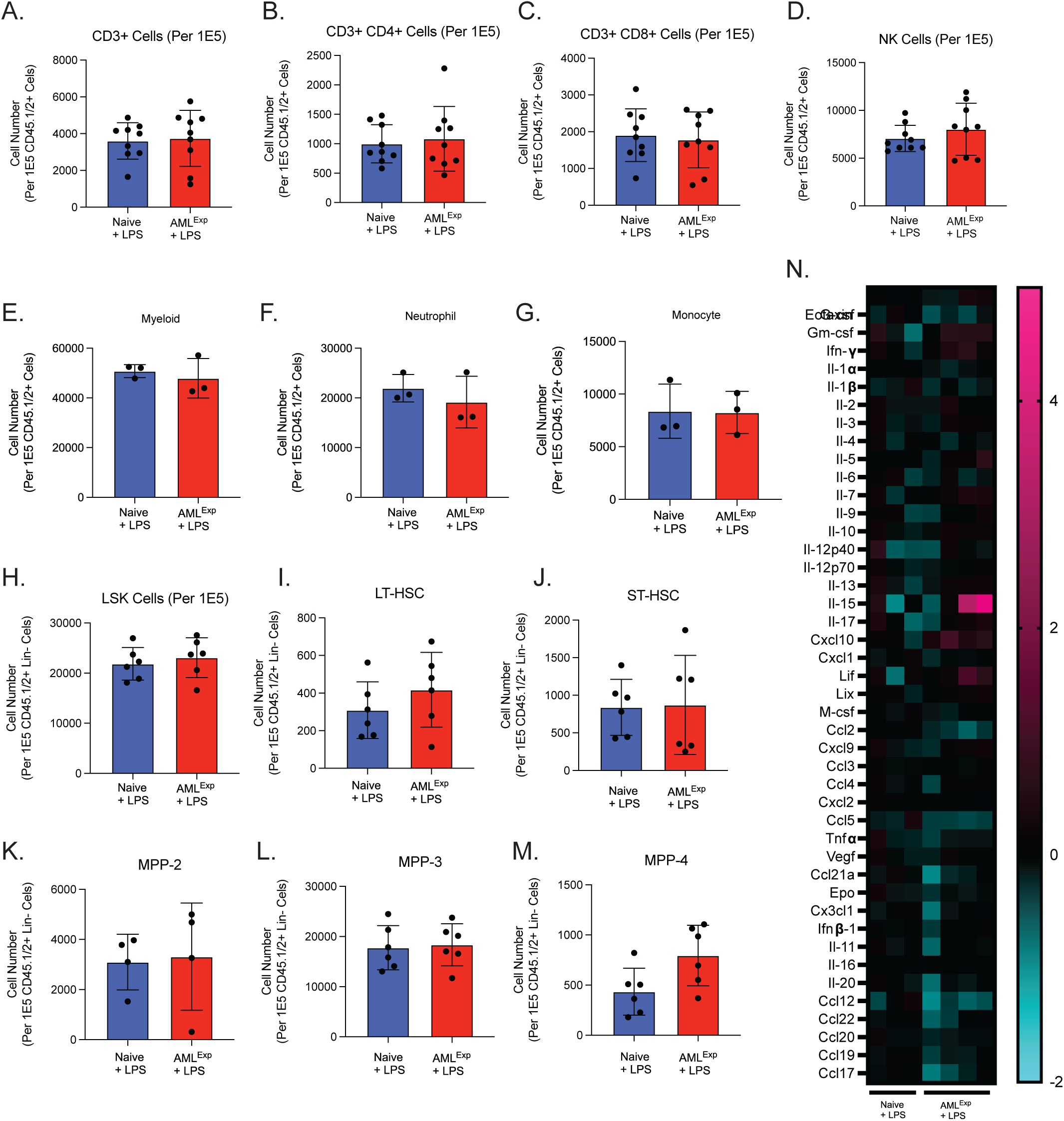
BM immune microenvironment characterization 24 hours after LPS challenge. Immunophenotype analysis of common mature lymphoid (A-D), myeloid (E-G), and SLAM population (H-M) in the BM (n=≥3). (N) 44-Plex chemokine/cytokine discovery analysis of BM plasma from LPS-challenged AML^EXP^ and Naïve mice (n=≥3). Values are expressed as mean ± standard deviation. Statistical significance was calculated using student’s t-test.

**Supplementary Figure 4.**
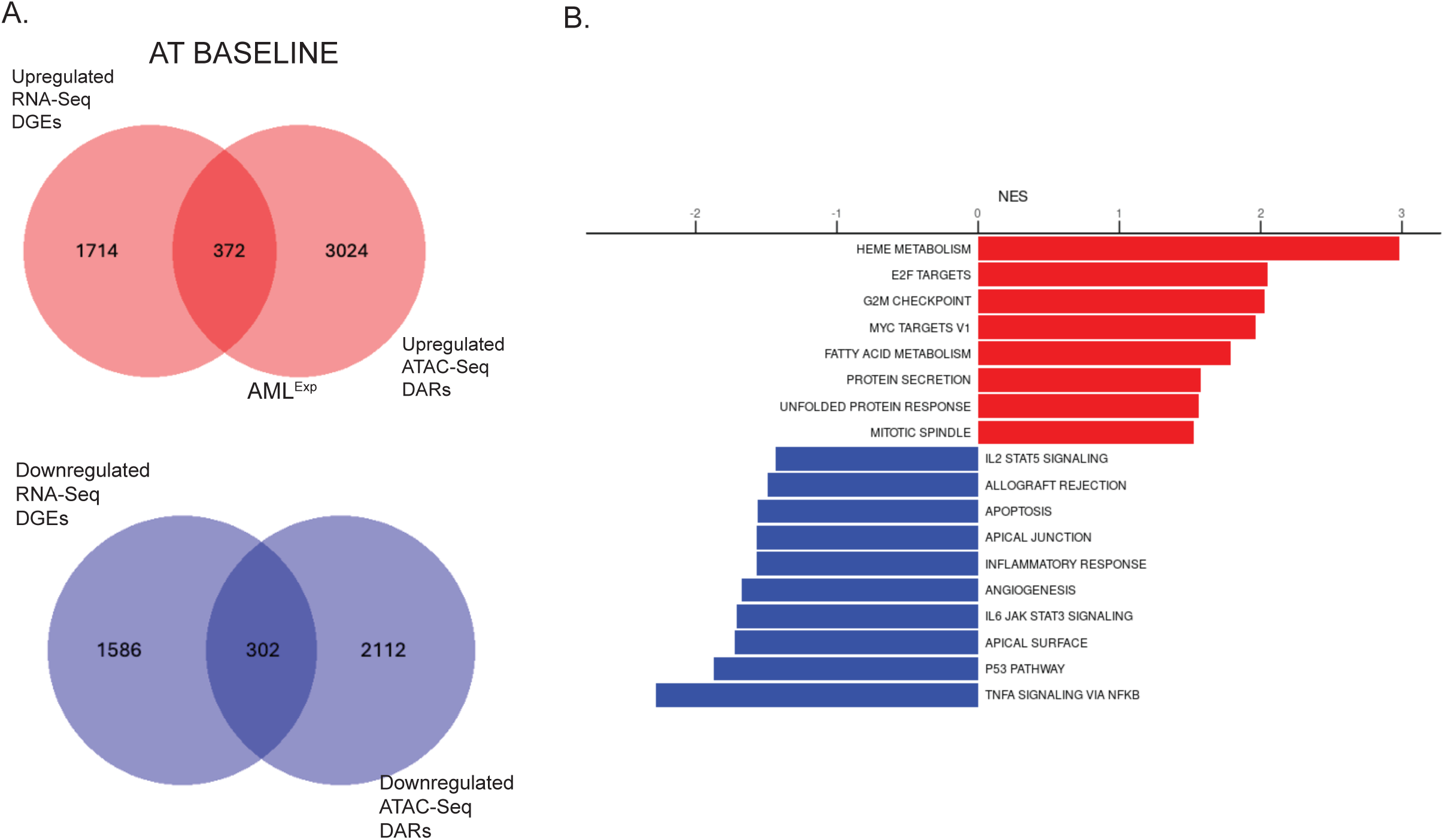
Correlative RNA-Seq and ATAC-Seq analysis. Correlative RNA-Seq and ATAC-Seq analysis in HSPC^AML+PBS^ and identification of genes that are differentially expressed and accessible (A). GSEA analysis of the common genes identified (B).

**Supplementary Figure 5.**
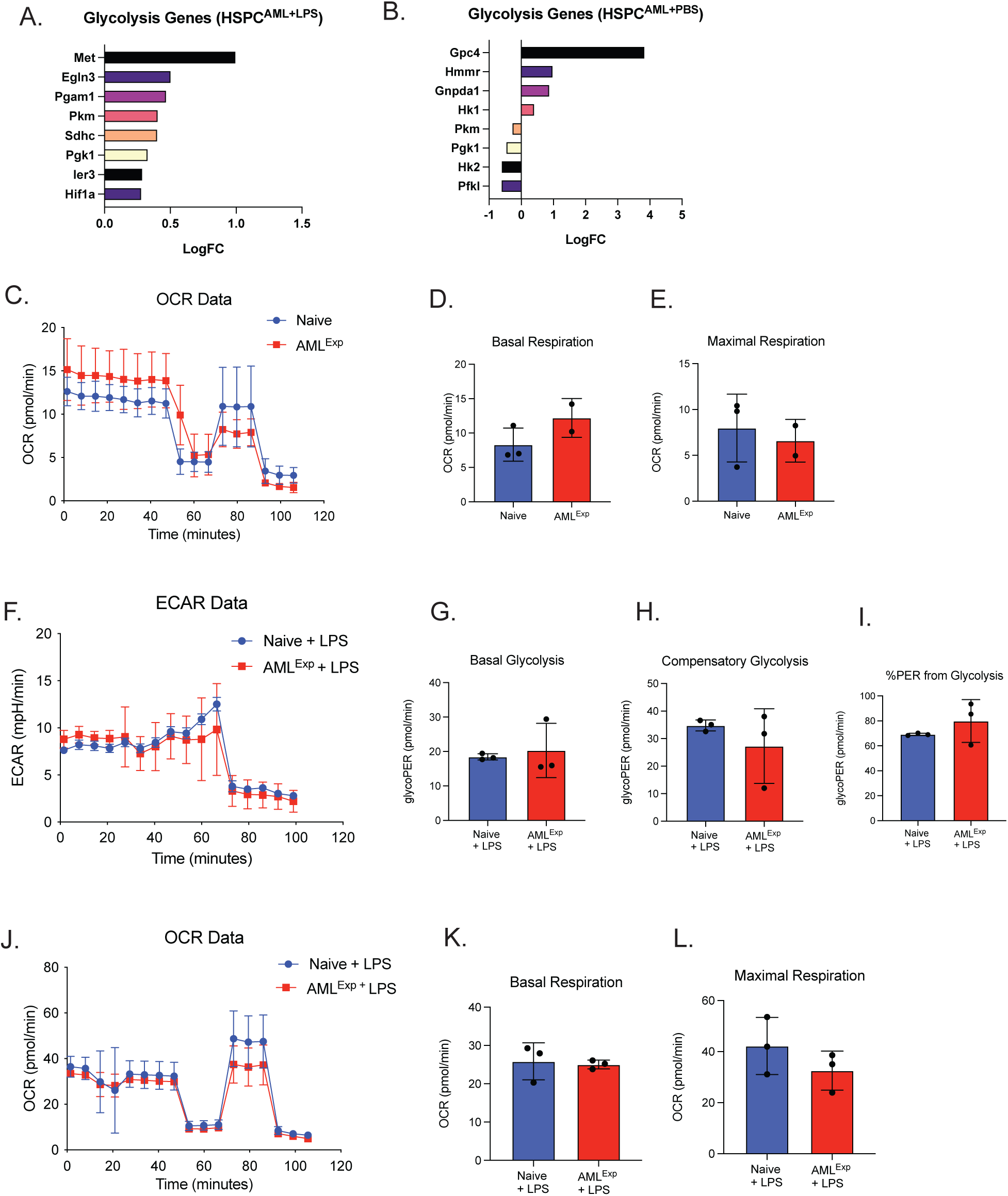
Glycolysis related gene expression in HSPC^AML+PBS^. (A) and HSPC^AML+LPS^ (B) (n=3). Mitochondrial function was assessed by subjecting HSPC^AML+PBS^ to Mito Stress assay (C-E) (n=3). HSPC^AML+LPS^ were also subjected to glycolytic rate assay (F-I) and mitochondrial stress assay (J-L). Values are expressed as mean ± standard deviation (s.d.), Statistical significance was calculated using student’s t-test. *p<0.05.

